# Reconfiguration of brain network architectures between resting state and complexity-dependent cognitive reasoning

**DOI:** 10.1101/163022

**Authors:** Luke J. Hearne, Luca Cocchi, Andrew Zalesky, Jason B. Mattingley

**Affiliations:** Queensland Brain Institute, The University of Queensland, Brisbane, Australia.; QIMR Berghofer Medical Research Institute, Brisbane, Australia.; Melbourne Neuropsychiatry Centre, University of Melbourne, Melbourne, Australia.; School of Psychology, The University of Queensland, Brisbane, Australia.

## Abstract

Our capacity for higher cognitive reasoning has a measureable limit. This limit is thought to arise from the brain’s capacity to flexibly reconfigure interactions between spatially distributed networks. Recent work, however, has suggested that reconfigurations of task-related networks are modest when compared with intrinsic ‘resting state’ network architecture. Here we combined resting state and task-driven functional magnetic resonance imaging to examine how flexible, task-specific reconfigurations associated with increasing reasoning demands are integrated within a stable intrinsic brain topology. Human participants (21 males and 28 females) underwent an initial resting state scan, followed by a cognitive reasoning task involving different levels of complexity, followed by a second resting state scan. The reasoning task required participants to deduce the identity of a missing element in a 4 × 4 matrix, and item difficulty was scaled parametrically as determined by relational complexity theory. Analyses revealed that external task engagement was characterized by a significant change in functional brain modules. Specifically, resting state and null-task demand conditions were associated with more segregated brain network topology, whereas increases in reasoning complexity resulted in merging of resting state modules. Further increments in task complexity did not change the established modular architecture, but impacted selective patterns of connectivity between fronto-parietal, subcortical, cingulo-opercular and default-mode networks. Larger increases in network efficiency within the newly established task modules were associated with higher reasoning accuracy. Our results shed light on the network architectures that underlie external task engagement, and highlight selective changes in brain connectivity supporting increases in task complexity.

**Significance Statement:** Humans have clear limits in their ability to solve complex reasoning problems. It is thought that such limitations arise from flexible, moment-to-moment reconfigurations of functional brain networks. It is less clear how such task-driven adaptive changes in connectivity relate to stable, intrinsic networks of the brain and behavioral performance. We found that increased reasoning demands rely on selective patterns of connectivity within cortical networks that emerged in addition to a more general, task-induced modular architecture. This task-driven architecture reverted to a more segregated resting state architecture both immediately before and after the task. These findings reveal how flexibility in human brain networks is integral to achieving successful reasoning performance across different levels of cognitive demand.

## Introduction

Humans are unparalleled in their ability to reason and solve complex problems in the service of goal-directed behavior (Penn et al., 2008; Johnson-Laird, 2010). Nevertheless, our ability to reason successfully is limited by the complexity of the task at hand (Halford et al., 1998, 2005). Increasing reasoning demands are supported by the flexible reconfiguration of large-scale functional brain networks (Cocchi et al., 2013, 2014), but recent work has demonstrated that such reconfigurations are relatively modest and occur within a preserved global network architecture (Cole et al., 2014; Krienen et al., 2014). Here we assessed changes in functional brain architecture induced by engagement in a complex reasoning task, as well as changes in communication across regions with parametric increases in reasoning complexity. To do so, we used high-field functional magnetic resonance imaging (fMRI) to measure brain activity at rest, and during performance of a behavioral task in which task complexity was manipulated parametrically.

Higher cognitive functions are supported by the adaptive reconfiguration of large-scale functional networks (Bassett et al., 2011; Cole et al., 2013; Braun et al., 2015; Cohen and D’Esposito, 2016; Yue et al., 2017). Previous empirical and theoretical work suggests that a multitude of complex tasks are related to activity and communication within and between select fronto-parietal, cingulo-opercular, and default-mode networks (Knowlton et al., 2012; Cocchi et al., 2014; Hearne et al., 2015; Crittenden et al., 2016; Bolt et al., 2017). Such networks are flexible and tend to increase their functional relationship in line with task demands across a wide range of domains, including reasoning (Cocchi et al., 2014), working memory (Vatansever et al., 2017) and decision making (Cole et al., 2013).

Recent empirical work has shown that task-induced network reconfigurations are modest when compared with intrinsic, ‘resting state’ networks (Cole et al., 2014; Krienen et al., 2014). For example, Cole and colleagues reported a matrix-level correlation between rest and task states of *r* = 0.90 (on average 38% of connections demonstrated change, with an average change of *r* = 0.04). Likewise, it is now apparent that task-induced activity can be well predicted and modeled from resting state data alone (Cole et al., 2016; Tavor et al., 2016). These results suggest that while behaviorally meaningful, selective task-induced reconfigurations occur against a backdrop of stable, large-scale networks that support diverse cognitive functions (Power et al., 2011; Crossley et al., 2013). An important unresolved question is how selective, ‘flexible’ task driven reconfigurations emerge amongst ‘stable’ intrinsic brain topology. Moreover, it is critical to understand how such global and selective changes are related to behavior (Bolt et al., 2017; Mill et al., 2017).

To investigate this question we measured functional brain networks at rest, as well as during several discrete levels of reasoning complexity. To systematically manipulate task complexity, we exploited *relational complexity theory* (Halford et al., 1998), which posits that the number of *relations* between variables quantifies the complexity of a problem, regardless of the domain of the original stimulus (e.g., semantic, spatial, etc.). Using this theoretical framework it has been shown that increasing the number of relations imposes a quantifiable cognitive load (measured via reaction time and accuracy), and eventually results in a breakdown of the reasoning process (Halford et al., 2005). We collected 7T fMRI data from 65 individuals while they undertook a non-verbal reasoning task known as the Latin Square Task (Birney et al., 2006). During the task, participants solved problems with three discrete levels of difficulty, defined formally in terms of their relational complexity (Binary, Ternary, Quaternary). In addition, just prior to the task, and again immediately afterwards, participants underwent a resting state scan. To examine network reconfigurations across rest and reasoning states we utilized *modularity* to assess segregation and integration and *global efficiency* to assess changes in network communication. Further to examine selective changes, we employed the network-based statistic to identify circumscribed changes in connectivity patterns (Zalesky et al., 2010), and related such network metrics to behavior.

## Materials and Methods

### Participants

Sixty-five healthy, right-handed participants undertook the current study, of whom 49 were included in the final analysis (*M* = 23.35 years, *SD* = 3.6 years, range = 18 – 33 years, 28 females). Four participants were excluded due to MR scanning issues, one participant was excluded due to an unforeseen brain structure abnormality, a further participant was excluded due to low accuracy in the behavioral task (total score more than 3 standard deviations below the mean) and ten participants were excluded due to excessive head movement (see *Preprocessing* section for head movement exclusions). Participants provided informed written consent to participate in the study. The research was approved by The University of Queensland Human Research Ethics Committee.

### Experimental paradigm

Each participant completed two behavioral sessions and one imaging session. In the imaging session participants underwent a resting state scan, followed by three, 12-minute runs of the Latin Square Task (LST; described below), a structural scan and finally a second resting state scan (see **Figure 1a**).

**Figure 1.**
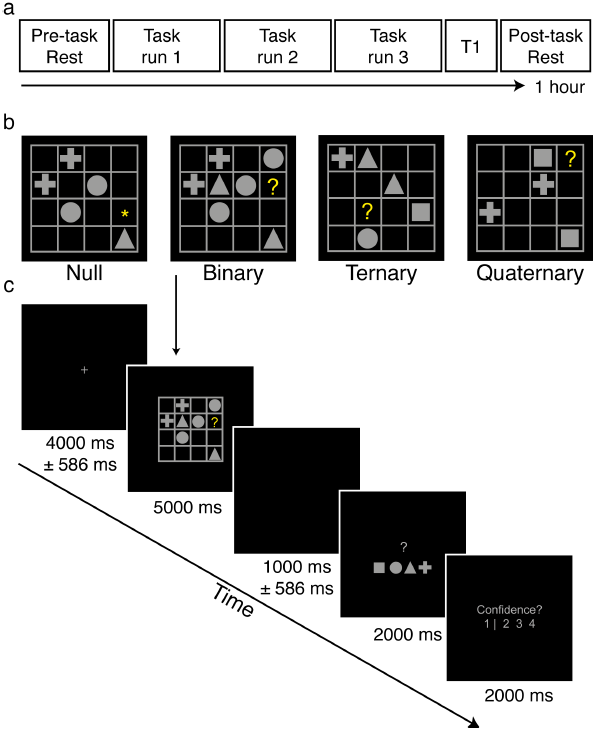
Experimental design and sequence of displays in a typical trial of the Latin Square Task. **a.** Functional magnetic resonance imaging session outline. Participants completed resting state scans before and after three runs of task imaging. **b.** Examples of each reasoning complexity condition. The correct answers are square, cross and cross, respectively, for the Binary, Ternary and Quaternary problems illustrated. **c.** Example trial sequence. Each trial contained a jittered fixation period, followed by an LST item, a second, jittered fixation period, a response screen, and a confidence rating scale. In Null trials the motor response screen had one geometric shape replaced with an asterisk, representing the correct button to press.

In the two behavioral sessions, participants completed the Raven’s Advanced Progressive Matrices (40 minute time limit), which is a standard and widely used measure of fluid intelligence (Raven, 2000). Of the 49 participants included in the analysis, 43 also completed a conjunction visual search task in which they were instructed to report the orientation of a target letter ‘L’ (rotated 90 degrees leftward or rightward) amongst ‘T’ distractors in set sizes of 8, 16 or 24 items. The search cost was defined as the increase in reaction time between the smallest and largest set sizes. This task was chosen as a ‘low reasoning’ counterpart to the Raven’s Progressive Matrices in order to demonstrate the specificity of brain-behavior correlations, as described in detail in the Results.

Participants also completed a modified version of the LST (Birney et al., 2006; LST, Birney and Bowman, 2009). The LST is a non-verbal relational reasoning paradigm in which reasoning complexity is parametrically varied with minimal working memory demands (Halford, 1998, Birney et al., 2006). Each LST ‘puzzle’ involves the presentation of a four-by-four matrix populated with a small number of geometric shapes (square, circle, triangle or cross), blank spaces and a single target, denoted by a yellow question mark (‘?’; see **Figure 1b**). Participants were asked to solve for the target according to the rule that *each shape can only occur once in every row and once in every column* (similar to the game of *Sudoku*). Binary problems require integration of information across a single row *or* column. Ternary problems involve integration across a single row *and* column. Quaternary problems, the most complex, require integration of information across multiple rows and columns (see **Figure 1b** for examples of each of these problems). Null trials involved presentation of an LST grid, but instead of a target question mark (‘?’) an asterisk was presented (‘*’) to cue the participant that no reasoning was required in this puzzle. The identity of the shapes that appeared in Null trials was random, but the number of shapes and their spatial locations were matched to those in the active LST trials. In total, 144 LST items were presented in the MR session across 16 blocks, with 36 items in each relational complexity condition; Null, Binary, Ternary and Quaternary. Prior to the MR session participants completed 20 practice trials of the LST (12 with corrective feedback). The visual angle subtended by the LST matrices was ~7.7 degrees, so that the entire stimulus fell within the parafoveal region of the visual field. Stimuli were projected onto a screen located at the head end of the MR scanner, and participants viewed the projected stimuli via a mirror mounted on the head coil.

Administration of all items was pseudo-randomized such that no two items of the same complexity occurred sequentially, and each block had two problems from each level of complexity (see **Figure 1c** for trial structure). Motor responses were counterbalanced across individuals, such that equal numbers of participants had the same shape-response mapping. Confidence ratings were used to determine participants’ subjective feeling of success, and to identify any trials in which participants inadvertently disengaged from the task altogether (e.g., due to a momentary lapse of attention). A three-point confidence scale indicated whether participants felt certain the problem had been answered correctly (4), felt unsure of their accuracy (3), or felt certain the problem had been answered incorrectly (2). On the far left (demarcated by a vertical line, see **Figure 1c**) was an additional ‘inattention’ rating point (1) that participants were instructed to select if they felt they had not attempted to solve the problem due to a momentary lapse of attention, fatigue or other factors. This response was used to separate incorrect choices arising from failures in reasoning, from those due to non-specific ”off-task” mind wandering (Smallwood and Schooler, 2015).

### Neuroimaging acquisition and preprocessing

Imaging data were collected using a 7 Tesla Siemens MR scanner fitted with a 32-channel head coil, at the Centre for Advanced Imaging, The University of Queensland. For both resting state and task fMRI, whole brain echo-planar images were acquired using a multi-band sequence (acceleration factor of five; Moeller et al., 2010). In each of the two resting scans, 1050 volumes were collected (~10 minutes each). In the each of the three runs of the task, 1250 volumes were collected (~12 minutes each) with the following parameters: voxel size = 2 mm^3^, TR = 586 ms, TE = 23 ms, flip angle = 40°, FOV = 208 mm, 55 slices. Structural images were also collected to assist functional data preprocessing. These images were acquired using the following parameters: MP2RAGE sequence, voxel size = 0.75 mm^3^, TR = 4300 ms, TE = 3.44 ms, 256 slices (see **Figure 1a** for session structure).

Imaging data were preprocessed using an adapted version of the MATLAB (MathWorks, USA) toolbox *Data Processing Assistant for Resting-State fMRI* (DPARSF V 3.0, Chao-Gan and Yu-Feng, 2010). Both resting state and task data were preprocessed with the same pipeline (except where noted). DICOM images were first converted to Nifti format and realigned. T1 images were re-oriented, skull-stripped (FSL BET), and co-registered to the Nifti functional images using statistical parametric mapping (SPM8) functions. Segmentation and the DARTEL algorithm were used to improve the estimation of non-neural signal in subject space and the spatial normalization (Ashburner, 2007). From each gray matter voxel the following signals were regressed: undesired linear trends, signals from the six head motion parameters (three translation, three rotation), white matter and cerebrospinal fluid (estimated from single-subject masks of white matter and cerebrospinal fluid). The CompCor method (Behzadi et al., 2007) was used to regress out residual signal unrelated to neural activity (i.e., five principal components derived from noise regions-of-interest in which the time series data were unlikely to be modulated by neural activity). Global signal regression was not performed due to the ongoing controversy associated with this step (Saad et al., 2012; Caballero-Gaudes and Reynolds, 2017). This choice may increase motion artifacts in the data (Ciric et al., 2017). For this reason we employed a strict head motion censoring approach (see below). Single-subject functional images were subsequently normalized and smoothed using DARTEL (4mm^3^). Data processing steps also involved filtering (0.01–0.15 Hz) at a low frequency component of the BOLD signal known to be sensitive to both resting state and task-based functional connectivity (Sun et al., 2004), therefore allowing comparison of both resting state and task data.

### Head movement

Participants with head displacement exceeding 3mm in more than 5% of volumes in any one scan were excluded. In addition to gross head movement, it has also been shown that functional connectivity can be influenced by small volume-to-volume ’micro’ head movements (Van Dijk et al., 2012; Power et al., 2014). To ensure micro-head movement artifacts did not contaminate our findings, both resting state and task-based data with frame-to-frame displacements greater than 0.40 mm were censored (Power et al., 2014). Participants with less than 85% of data remaining in any condition were excluded.

### Functional connectivity network construction

For each subject, regionally averaged time series were extracted for 264 spheres of 5mm radius sampled across cortical and subcortical gray matter. Spheres were positioned according to an existing brain parcellation, based on task activations induced by a wide range of behavioral tasks (Power et al., 2011). This parcellation and associated network definitions were generated from a large cohort of participants (N > 300), and has the advantage of being independent of the imaging data obtained in the current study.

For both sets of resting state data (pre-and post-task), functional connectivity was estimated using a temporal Pearson correlation between each pair of time series (Zalesky et al., 2012). This resulted in a 264 × 264 connectivity matrix for each subject. For the task-based functional connectivity analyses we used a redgression approach (i.e., Cole et al., 2014) rather than psycho-physiological interactions (PPI) as others have used previously (McLaren et al., 2012; Cocchi et al., 2014; Gerchen et al., 2014). We opted for this approach rather than PPI due to our interest in assessing connectivity across both rest and task states. For each brain region of interest, a task regressor composed of the condition onsets modeled as boxcar functions convolved with a canonical hemodynamic response function was regressed from the time series. This step was taken to remove variance associated with task-related coactivation (Cole et al., 2014). Then, after accounting for the hemodynamic lag, the residual time series from each five-second reasoning period was concatenated to form a condition-specific time series of interest, in each brain region. A Pearson correlation was performed on the resulting regional time series for each condition separately resulting in a 4 (condition) x 264 ×264 connectivity matrix for each subject. Finally, both resting state and task-based matrices were converted to z-scores. Analysis decisions such as z-normalization and thresholding were employed so as to be consistent with previous, related work aimed at assessing dynamic reconfiguration of connectivity patterns as a function of task demands (e.g., Cole et al., 2014, Power et al., 2011). Such choices do, however, affect the resulting graph metrics (Rubinov and Sporns, 2011). Thus, unless otherwise noted (see network based statistic analysis, below) weighted graphs of proportional densities from the top 5% to the top 30% of connections were considered for analysis. Such network densities have been shown to provide robust functional brain network characterizations (Garrison et al., 2015) and are similar to those used in previous, related work (e.g., Power et al., 2011).

### Analysis overview

We undertook three complementary analyses to identify functional network reorganization due to increasing relational complexity. First, we calculated and compared community partitions that arose in each of the resting state and task conditions. Following this, we performed an analysis to identify changes in connectivity associated with performance of the Latin Square Task using the network based statistic (NBS, Zalesky et al., 2010), a sensitive statistical tool that controls for Type I error at the network level. To assess the functional and behavioral impact of the connectivity changes identified in the previous two analyses, we calculated changes in *global efficiency* (Achard and Bullmore, 2007) for each functional module detected. Moreover, to assess the behavioral implications of the observed network changes, we correlated metrics of changes in module efficiency with performance accuracy on the LST. When appropriate, nonparametric statistics were used for repeated-measures comparisons (Friedman test), follow-up tests (Wilcoxon signed rank) and measures of effect size (Kendall’s coefficient of concordance, *W*).

### Community detection

A module is a group of nodes in a graph that contains stronger connections within-module than expected in an appropriate random network null model. A modularity partition represents the subdivision of a graph into non-overlapping modules (Fortunato, 2010). The degree of modularity in a network can be characterized by the Q index (Newman and Girvan, 2004), which represents the density of within-module connections relative to an appropriate random network null model. The aim of community detection is to isolate a module partition that maximizes Q.

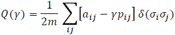

Above is the modularity equation, where a_*ij*_ represents the weight of the edge between *i* and *j*, 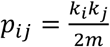 represents the expected number of links according to the so-called configuration null model (Newman et al., 2001), where *k*_*i*_ is the degree of node *i*; in this case the null preserves the node degree while forming connections at random. *2m* represents the total number of connections in the network, σ_*i*_ denotes the community to which node *i* is assigned, the Kronecker delta function, δ(σ_*i*_σ_*j*_), is 1 if σ_*i*_ = σ_*j*_ and 0 if otherwise. Finally, γis the resolution parameter; when γ < 1, larger communities are resolved, if γ > 1, smaller communities are resolved.

In the present study, modules were identified using the Louvain greedy algorithm (Blondel et al., 2008) implemented in the Brain Connectivity Toolbox (BCT, Rubinov and Sporns, 2010). The resolution parameter was set to unity (γ = 1). Testing across several levels of γ showed consistent results. For clarity, we highlight the BCT scripts used throughout the Method section. There are multiple possible module partitions that maximize Q for each graph, resulting in community assignments that vary across each run of the algorithm (Good et al., 2010; Sporns and Betzel, 2016). To resolve this variability we used a consensus approach (Lancichinetti and Fortunato, 2012), whereby module partitions are calculated a number of times (10^3^ iterations, for each participant and condition) and used to calculate an agreement matrix (agreement.m). The agreement matrix represents the tendency for each pair of nodes to be assigned to the same module across iterations. Finally, the agreement matrix was subjected to an independent module partitioning (consensus_und.m), resulting in an individual-level module partition for each participant in each condition. In this step, the resolution parameter was also set to unity (τ = 1), representing the level at which the agreement matrix was thresholded before being subjected to the consensus procedure. For example, τ = 1 thresholds the matrix such that only nodes consistently partitioned into the same community across all permutations are included. Testing across several levels of τ showed consistent results. A general community structure including motor-sensory, auditory, visual, default-mode and fronto-parietal/cingulo-opercular modules was entered into the algorithm as the initial community partition. In our data, this choice decreased computation time, presumably because the initial community structure was associated with a Q-value that was close to the true maximum. Module reconfiguration results were replicated using different community affiliation priors, including variations of the original community partitions (Power et al., 2011; Cole et al., 2013) and purely data-driven methods (i.e., no community affiliation input).

The procedure for group-level modular decomposition was implemented in a similar fashion to the individual-level decompositions described above. The critical difference was that instead of creating an individual-level agreement matrix, the agreement matrix represented the tendency for each pair of nodes to be assigned to the same module across participants. The same consensus procedure followed, resulting in a single module partition for each condition for the group of 49 participants. Resolution parameters were kept identical to the previous individual-level modularity analysis.

### Significance testing for within-participant differences in modular structure

To investigate differences in the nodal composition of modules across conditions, we used the Variation of Information metric (*VIn*, Meilă, 2007), an information-theoretic measure of partition distance (*partition_distance.m*). To ascribe statistical significance to differences in partition structure we used a repeated-measures permutation procedure to compare real VIn values to appropriate null distributions (Dwyer et al., 2014). Specifically, half of the participants’ condition labels were randomly switched in the contrast of interest (e.g., Binary versus Ternary). This resulted in two new sets of individual-level module structures for the contrast (albeit with shuffled data). The shuffled module structures were then subjected to the previously used pipeline to generate group-level module partitions. Finally, *VIn* was used to quantify the difference between these partitions. This procedure was repeated 10^4^ times to build a null distribution for each contrast of interest, with which the real data were compared.

### Pairwise functional connectivity analysis

The Network Based Statistic (NBS, Zalesky et al., 2010) was used to identify changes in pairwise functional connectivity at rest and during the task. For the first contrast, a paired *t*-test was performed between the Pre-and Post-task resting state data. For the second contrast, a one-way repeated measures ANOVA was used to compare all four task states (Null, Binary, Ternary, Quaternary). For the analysis, *unthresholded* functional connectivity matrices were used as input into the NBS. Briefly, all possible pairs of connections (264 × 263/2 = 34,716) were tested against the null hypothesis, endowing each connection with a test statistic, which was subsequently thresholded. Here an exploratory F-statistic of 20 (equivalent to a *t*-statistic of 4.47) was used as the threshold, though additional exploratory analyses showed that networks arising using higher or lower t-thresholds resembled the original results. This threshold was adopted because it allowed the detection of effects of medium size while discarding small or spurious effects. Family-wise error corrected (FWE) *p*-values were ascribed to the resulting networks using a null distribution obtained by 5,000 permutations. Only components that survived a network-level threshold of *p* < 0.001 FWE were declared significant. This analysis allowed us to identify sub-networks that significantly *increased* or *decreased* their functional connectivity across relational reasoning task conditions, providing complementary results to the graph analyses.

### Network efficiency analysis

Global efficiency is defined as the inverse of the average characteristic path length between all nodes in a network (Latora and Marchiori, 2001). Assuming that information follows the most direct path, global efficiency provides an index for parallel information transfer in a network (Rubinov and Sporns, 2010). In the context of functional brain networks, global efficiency is thought to be an index of increased capacity for information exchange (Achard and Bullmore, 2007). The link between indices of global efficiency and global neural information transfer is, however, not yet clear. Nevertheless, a number of studies have shown that high global brain network efficiency can enhance neurophysiological (De Pasquale et al., 2016; Cocchi et al., 2017) and cognitive processes (Bassett et al., 2009; van den Heuvel et al., 2009; Shine et al., 2016).

Here we wanted to investigate differences in network communication *within module,* and determine how such difference might relate to behavior. To do so, we computed global efficiency for each participant, in each condition, for the three major modules identified in the initial modularity analysis (using *efficiency_wei.m* from the BCT). Importantly, matrix thresholding was performed after dividing the modules to ensure any efficiency effects were not due to differences in degree across modules. Finally, we computed the difference in efficiency between the most and least difficult conditions (i.e., Quaternary versus Null) and correlated this change in efficiency with overall accuracy scores on the Latin Square Task.

### Figures and Visualization

Figures were generated with a combination of MATLAB, and online network visualization tools (*alluvial* diagram; (http://www.mapequation.org/apps/MapGenerator.html, and the *connectogram*; http://immersive.erc.monash.edu.au/neuromarvl/).

## Results

### Behavioral results

A nonparametric Friedman test revealed a significant effect of reasoning complexity on both LST accuracy (χ^2^ = 86.20, Kendall’s *W* = 0.88, *p* < 0.001) and reaction time (χ^2^ = 63.71, *W* = 0.65, *p* < 0.001, see **Figure 2**). Bonferroni corrected follow-up Wilcoxon signed-rank test comparisons revealed that accuracy was significantly higher for the Binary condition (*M* = 34.96, *SD* = 1.04) than for both the Ternary condition (*M* = 31.63, *SD* = 3.52, *z* = 5.59, *p* < 0.001) and the Quaternary condition (*M* = 22.78, *SD* = 6.71, *z* = 6.10, *p* < 0.001). Accuracy was also higher for Ternary items than for Quaternary items (*z* = 5.74, *p* < 0.001). The reaction time results followed a similar pattern, such that responses were faster in the Binary condition (*M* = 771.60 ms, *SD* = 201.60 ms) than in the Ternary (*M* = 844.20 ms, *SD* = 217.30 ms, *z* = -4.62, *p* < 0.001) and Quaternary (*M* = 933.70 ms, *SD* = 205.00 ms, *z* = -5.83, *p* < 0.001) conditions. Likewise, reaction times in the Ternary condition were significantly faster than those in the Quaternary condition (*z* = -4.79, *p* < 0.001).

**Figure 2.**
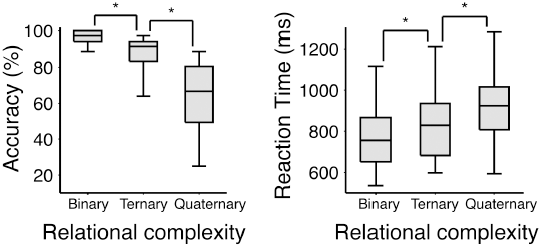
Behavioral results for the Latin Square Task visualized as box and whisker plots. Here the boxes represent the median and interquartile ranges, and the whiskers show the minimum and maximum values. **a**. Accuracy as a function of reasoning complexity. **b.** Reaction time as a function of reasoning complexity. Significance markers indicate p < 0.001.

As expected, there was a significant positive correlation between scores on the Raven’s Advanced Progressive Matrices, a measure of fluid intelligence, and overall accuracy on the LST (*r* = 0.44, *p* = 0.002). By contrast, for the visual search task, there was no correlation between reaction time cost and LST score (*N* = 43, *r* = -0.09, *p* = 0.58). A Steiger *z*-test (Lee and Preacher, 2013) demonstrated that these two correlations were significantly different from one another (*N* = 43, *z* = -2.90, *p* = 0.003), confirming that LST performance is linearly related to an established measure of fluid intelligence, but not to a widely used test of visual attention (Triesman & Gelade, 1980).

Participants’ confidence was assessed on each trial. Importantly, we included an explicit rating for when participants had not attempted the reasoning problem due to an attention lapse, mind wandering, or fatigue. Averaging across all conditions, the mean number of such lapses was fewer than one out of 36 trials (*M* across conditions = 0.47 trials, *SD* across conditions = 0.98 trials). We can thus conclude that overall, participants were able to engage as instructed in cognitive reasoning across all three levels of relational complexity in the LST.

### Functional brain module reconfiguration

Modularity analysis revealed four major modules in the baseline (Pre-task) resting state. For clarity, these modules are represented in reference to Power and colleagues’ (2011) initial network affiliations. The modules broadly correspond to the *sensory, default-mode, visual* and *fronto-parietal* networks.

Variation of Information analysis (Meilă, 2007) revealed a significant difference in the community structure of Binary, Ternary and Quaternary conditions compared with the Pre-task resting state (mean statistics across thresholds are reported in text; see **Table 1** for extensive results, *VIn* = 0.20, *p* = 0.006, *VIn* = 0.21, *p* = 0.001, *VIn* = 0.20, *p* = 0.003, respectively) and Post-task resting state (*VIn* = 0.18, *p* = 0.039, *VIn* = 0.20, *p* = 0.005, *VIn* = 0.19, *p* = 0.016). The difference between rest and reasoning states was associated with the emergence of a single, conjoined *fronto-parietal-visual* module that was composed of several large-scale networks identified by Power et al. (2011). The transitory nature of the *fronto-parietal-visual* module was confirmed by its switch back to its original configuration in the Post-task resting state (i.e., after completion of the LST). **Figure 3b** shows a representation of the reconfiguration of modules across experimental conditions. There was no significant difference between the Pre-task and Post-task resting state community structure (*VIn* = 0.09, *p* = 0.788). There was also no consistent difference between the Pre-and Post-task resting state communities and the Null task condition (*VIn* = 0.16, *p* = 0.08, *VIn* = 0.15 *p* = 0.196, respectively). These main effects were broadly replicated across all thresholds tested (**Figure 3c**).

**Table 1.** Variation of Information statistics.

|  |  | Network density |  |  |  |  |  |  |  |  |  |  |  |
| --- | --- | --- | --- | --- | --- | --- | --- | --- | --- | --- | --- | --- | --- |
|  |  | 5% |  | 10% |  | 15% |  | 20% |  | 25% |  | 30% |  |
| Contrast |  | VIn | *p* | VIn | *p* | VIn | *p* | VIn | *p* | VIn | *p* | VIn | *p* |
| Pre-task rest | Post-task rest | 0.056 | 0.878 | 0.065 | 0.863 | 0.109 | 0.656 | 0.098 | 0.836 | 0.124 | 0.552 | 0.092 | 0.944 |
|  | Null | 0.146 | 0.013 | 0.139 | 0.077 | 0.124 | 0.397 | 0.189 | 0.009* | 0.198 | 0.005* | 0.186 | 0.029 |
|  | Binary | 0.148 | 0.006* | 0.179 | 0.001* | 0.218 | 0.001* | 0.219 | < 0.001* | 0.216 | 0.007* | 0.209 | 0.023 |
|  | Ternary | 0.166 | 0.002* | 0.194 | < 0.001* | 0.226 | < 0.001* | 0.226 | 0.001* | 0.221 | 0.002* | 0.215 | 0.002* |
|  | Quaternary | 0.148 | 0.009* | 0.189 | < 0.001* | 0.225 | < 0.001* | 0.22 | < 0.001* | 0.221 | < 0.001* | 0.203 | 0.008* |
| Post-task rest | Null | 0.117 | 0.231 | 0.154 | 0.019 | 0.098 | 0.81 | 0.153 | 0.084 | 0.185 | 0.015 | 0.191 | 0.018 |
|  | Binary | 0.128 | 0.099 | 0.185 | 0.001* | 0.176 | 0.033 | 0.181 | 0.074 | 0.212 | 0.01* | 0.203 | 0.018 |
|  | Ternary | 0.16 | 0.006* | 0.204 | < 0.001* | 0.194 | 0.006* | 0.203 | 0.003* | 0.2 | 0.016 | 0.22 | 0.001* |
|  | Quaternary | 0.128 | 0.083 | 0.189 | 0.002* | 0.195 | 0.005* | 0.202 | 0.001* | 0.2 | 0.005* | 0.211 | 0.001* |
| Null | Binary | 0.028 | 1 | 0.121 | 0.245 | 0.162 | 0.199 | 0.1 | 0.698 | 0.1 | 0.674 | 0.106 | 0.534 |
|  | Ternary | 0.094 | 0.711 | 0.133 | 0.237 | 0.193 | 0.023 | 0.13 | 0.645 | 0.119 | 0.567 | 0.132 | 0.246 |
|  | Quaternary | 0.052 | 0.987 | 0.117 | 0.272 | 0.181 | 0.04 | 0.14 | 0.539 | 0.132 | 0.467 | 0.132 | 0.291 |
| Binary | Ternary | 0.096 | 0.681 | 0.154 | 0.159 | 0.097 | 0.801 | 0.1 | 0.485 | 0.093 | 0.571 | 0.11 | 0.306 |
|  | Quaternary | 0.038 | 0.998 | 0.121 | 0.375 | 0.086 | 0.91 | 0.111 | 0.507 | 0.106 | 0.484 | 0.108 | 0.387 |
| Ternary | Quaternary | 0.082 | 0.747 | 0.084 | 0.894 | 0.041 | 1 | 0.051 | 0.986 | 0.061 | 0.943 | 0.051 | 0.969 |

Note: \* indicates contrasts where $p < 0.01$

**Figure 3.**
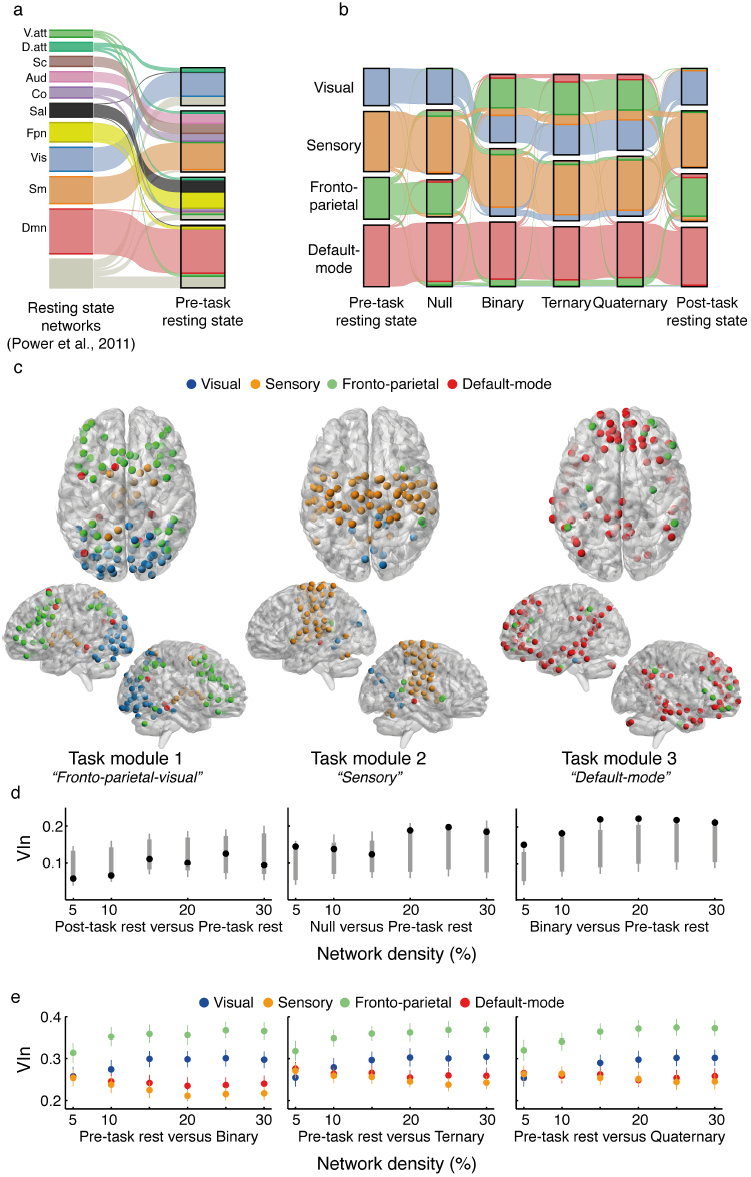
Modular structure as a function of reasoning complexity in the Latin Square Task (LST). **a.** Alluvial ‘flow’ demonstrating the network affiliations (as per Power et al., 2011) compared with the Pre-task resting state (Rosvall et al., 2009). Each individual streamline represents a node in the network, colored by its original resting state affiliation as shown on the left (Power et al., 2011). **b.** Changes in modular structure across the experimental conditions. Visual and fronto-parietal modules merged to form a ‘Task-related’ module during Binary, Ternary and Quaternary conditions of the LST. Results for 15% network density are shown, but statistics were performed across several thresholds. **c.** Anatomical rendering of the task-related modules in the Quaternary condition. Each sphere is color-coded by its initial resting state module allegiance. **d.** Variation of Information (VIn) values (black markers) compared with a null distribution (gray markers, 5^th^/95^th^ percentile in bold line, 1^st^/99^th^ percentile shown in tails) for the three main contrasts across all network densities. Only the right-most contrast (Binary versus Pre-task rest) showed a consistent difference between partitions. **e.** Comparison of VIn values across visual, sensory, fronto-parietal and default-mode modules in each task condition compared with rest across all network densities. The fronto-parietal module was consistently *more variable* in relation to other modules. Error bars represent 95% confidence intervals.

Having established a difference in community structure we sought to test the relative contribution of each module to the observed reconfiguration. To do so we implemented a similar strategy to Braun and colleagues (2015), whereby VIn was calculated at the individual-community level for our nodes of interest, and compared using repeated-measures statistics. Thus we compared visual, sensory, fronto-parietal and default-mode modules across all network densities for the Binary-Rest, Ternary-Rest, and Quaternary-Rest contrasts (conceptually similar to follow-up parametric statistics). Results revealed that the fronto-parietal module had higher VIn values (i.e., larger differences in community structure) than visual (mean *p* value across thresholds, *p* < 0.001), sensory (*p* < 0.001) and default-mode modules (*p* < 0.001), see **Figure 3d**) across all contrasts.

Finally, the index of modularity, Q, was compared across conditions. This index of modularity increases as more intramodular connections are found than expected by chance (Newman & Girvan, 2004). Non-parametric Friedman tests revealed a significant difference in Q across conditions (mean statistic across thresholds, χ^2^ = 51.48, *p* < 0.001). Bonferonni corrected follow-up tests were performed to compare each task state (Null, Binary, Ternary, Quaternary) with each resting state (Pre-and Post-task). Results revealed that Q was significantly lower in the Ternary (Mean Q = 0.39) and Quaternary conditions (Mean Q = 0.39) when compared with both Pre-(Mean Q = 0.44, *z* = 3.74, *z* = 4.07, *p* < 0.001) and Post-task resting states (Mean Q = 0.45, *z* = 4.49, *z* = 4.52, *p* < 0.001). No effect was found when comparing Null or Binary conditions with rest. Complementing the observed changes in community structure, analysis of Q scores highlight a significant reduction in modularity compared with the resting state, but only in the task conditions that imposed higher demands on cognitive reasoning.

#### Network Based Statistic analysis

To further refine our account of module reconfiguration, we assessed changes in whole brain connectivity using the network based statistic (NBS, Zalesky et al., 2010). In line with the result from the first analysis, a paired t-test between Pre-and Post-task resting states revealed no significant differences. Our second contrast, a one-way repeated measures ANOVA was performed comparing all four reasoning complexity conditions (Null, Binary, Ternary, Quaternary).

A sub-network comprising 63 nodes and 85 edges changed in response to reasoning complexity demands (*p* < 0.001, FWE corrected at the network level, **Figure 4**). The majority of edges within the subnetwork demonstrated increased functional connectivity (86% of edges, shown as warm colors in **Figure 4**), but a number of edges also demonstrated a decrease in *positive* correlations with increasing reasoning complexity (see **Figure 4b** for trend across conditions). Consistent with our previous work on changes in functional connectivity during complex reasoning (Cocchi et al., 2014; Hearne et al., 2015), the network was largely composed of nodes encompassing fronto-parietal (17%), subcortical (19%), cingulo-opercular (12%) and default-mode networks (24%, as per Power et al., 2011 network affiliations, see **Table 2** for a list of regions implicated by this analysis). Moreover, nearly all edges (95%) were across-network. Two further visual-parietal subnetworks were identified by the NBS, consisting of two and three nodes respectively (not visualized).

**Table 2.** Significant pairwise changes in functional connectivity associated with increasing relational complexity

| x | MNI |  | RSN | Mod | Anatomy | Degree |
| --- | --- | --- | --- | --- | --- | --- |
|  | y | z |  |  |  |  |
| -7 | -52 | 61 | SShand | Vis | Precuneus | 12 |
| 4 | -48 | 51 | Mem | Sens | Precuneus | 10 |
| 47 | 10 | 33 | FPN | FPN | Precentral gyrus | 7 |
| -2 | -35 | 31 | Mem | DMN | Middle cingulate cortex | 6 |
| -34 | 3 | 4 | CO | Vis | Middle insula | 6 |
| 23 | 10 | 1 | SC | Vis | Putamen | 6 |
| 49 | 8 | -1 | CO | FPN | Superior temporal pole | 5 |
| -11 | -56 | 16 | DMN | DMN | Calcarine gyrus | 5 |
| -2 | 38 | 36 | DMN | DMN | Medial superior frontal gyrus | 5 |
| -42 | -55 | 45 | FPN | FPN | Inferior parietal cortex | 5 |
| -45 | 0 | 9 | CO | Vis | Rolandic operculum | 4 |
| 52 | -59 | 36 | DMN | DMN | Angular gyrus | 4 |
| -35 | 20 | 51 | DMN | DMN | Middle frontal gyrus | 4 |
| 9 | -4 | 6 | SC | Vis | Thalamus | 4 |
| -2 | -13 | 12 | SC | Vis | Thalamus | 4 |
| -47 | 11 | 23 | FPN | FPN | Inferior frontal gyrus (operculum) | 4 |
| -42 | 45 | -2 | FPN | FPN | Inferior frontal gyrus (orbital) | 4 |
| 54 | -28 | 34 | CO | Vis | Supramarginal gyrus | 3 |
| 8 | -48 | 31 | DMN | DMN | Posterior cingulate cortex | 3 |
| 13 | 30 | 59 | DMN | DMN | Superior frontal gyrus | 3 |
| 11 | -39 | 50 | Sal | Sens | Middle cingulate cortex | 3 |
| 15 | -77 | 31 | Vis | Sens | Cuneus | 3 |
| -41 | 6 | 33 | FPN | FPN | Precentral gyrus | 3 |
| -53 | -49 | 43 | FPN | FPN | Inferior parietal cortex | 3 |
| -44 | 2 | 46 | FPN | DMN | Precentral gyrus | 3 |
| 3 | -17 | 58 | SShand | Vis | Supplementary motor area | 2 |
| 65 | -33 | 20 | Aud | Vis | Superior temporal gyrus | 2 |
| -51 | 8 | -2 | CO | Vis | Insula | 2 |
| 15 | -63 | 26 | DMN | DMN | Cuneus | 2 |
| 11 | -54 | 17 | DMN | DMN | Precuneus | 2 |
| 36 | 10 | 1 | CO | FPN | Insula | 2 |
| -10 | -18 | 7 | SC | Vis | Thalamus | 2 |
| -22 | 7 | -5 | SC | Vis | Putamen | 2 |
| 43 | -78 | -12 | Vis | Sens | Inferior occipital cortex | 2 |
| -42 | -60 | -9 | DorAtt | Sens | Inferior temporal gyrus | 2 |
| -3 | 26 | 44 | FPN | FPN | Superior frontal gyrus (medial) | 2 |
| 44 | -53 | 47 | FPN | FPN | Inferior parietal cortex | 2 |
| 32 | 14 | 56 | FPN | DMN | Middle frontal gyrus | 2 |
| -40 | -19 | 54 | SShand | Vis | Precentral gyrus | 1 |
| 0 | -15 | 47 | SShand | Vis | Middle cingulate cortex | 1 |
| -10 | -2 | 42 | CO | Vis | Middle cingulate cortex | 1 |
| -50 | -34 | 26 | Aud | Vis | Supramarginal gyrus | 1 |
| 59 | -17 | 29 | Aud | Vis | Postcentral gyrus | 1 |
| 37 | 1 | -4 | CO | Vis | Insula | 1 |
| 6 | -59 | 35 | DMN | DMN | Precuneus | 1 |
| -41 | -75 | 26 | DMN | DMN | Middle occipital cortex | 1 |
| 65 | -31 | -9 | DMN | DMN | Middle temporal gyrus | 1 |
| 43 | -72 | 28 | DMN | FPN | Middle occipital cortex | 1 |
| -3 | 42 | 16 | DMN | DMN | Anterior cingulate cortex | 1 |
| -16 | 29 | 53 | DMN | DMN | Superior frontal gyrus | 1 |
| 2 | -24 | 30 | Mem | DMN | Middle cingulate cortex | 1 |
| 12 | 36 | 20 | DMN | DMN | Anterior cingulate cortex | 1 |
| 6 | -24 | 0 | SC | Vis | Thalamus | 1 |
| 12 | -17 | 8 | SC | Vis | Thalamus | 1 |
| -5 | -28 | -4 | SC | Vis | Lingual gyrus | 1 |
| 31 | -14 | 2 | SC | Vis | Putamen | 1 |
| 29 | 1 | 4 | SC | Vis | Putamen | 1 |
| -31 | -11 | 0 | SC | Vis | Putamen | 1 |
| 15 | 5 | 7 | SC | Vis | Pallidum | 1 |
| 37 | -84 | 13 | Vis | Sens | Middle occipital cortex | 1 |
| 37 | -81 | 1 | Vis | Sens | Middle occipital cortex | 1 |
| -33 | -79 | -13 | Vis | Sens | Inferior occipital cortex | 1 |
| 49 | -42 | 45 | FPN | FPN | Inferior parietal cortex | 1 |

Note: Automated Anatomical Labeling atlas was used to define anatomical regions. RSN = initial Power et al., 2011 resting state network affiliation. Mod = modules defined by modularity analysis (i.e., **Figure 3**).

**Figure 4.**
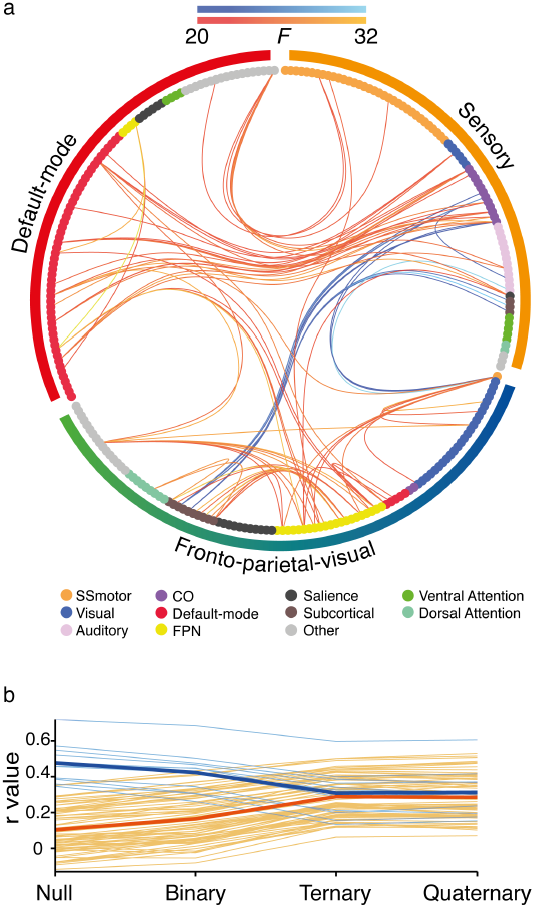
Change in pairwise functional connectivity associated with reasoning complexity. **a.** Connectogram representation of significant changes in pairwise functional connectivity that scaled with relational complexity. Edges are colored by the direction of the change in correlation across relational complexity. Warm colors represent increases in connectivity and cool colors represent decreases in connectivity. Lighter colors represent higher *F*-statistics. Network nodes, plotted as circles, are colored by their initial resting state networks (Power et al., 2011). Outside the connectogram the colored bars represent the modules identified in the previous analysis of data from the Quaternary condition: *Sensory* (orange), *Default-mode* (red) and *Fronto-parietal-visual* modules (green-blue). **b.** Each individual connection in the subnetwork (averaged across subjects) plotted as a function of reasoning complexity. Average values for positive and negative connections are shown as bold lines.

#### Within-module global efficiency and behavior

Our final analysis sought to investigate changes in global efficiency within each major module evident during the task. Global efficiency has previously been taken to be an index of increased capacity for information exchange (Achard and Bullmore, 2007). Specifically, we were interested in whether each module showed changes in efficiency, and whether any such changes were related to reasoning performance.

Nonparametric Friedman tests revealed that both the Sensory (orange in **Figure 5a**) and Fronto-parietal-visual modules (FPV, green in **Figure 5a**) demonstrated significant differences in efficiency across conditions (Sensory χ^2^ = 44.17, *p* < 0.001, FPV: χ^2^ = 77.78, *p* < 0.001, see **Figure 5a**). No such effect was found for the Default-mode module (*p* = 0.23). Bonferroni corrected follow-up tests confirmed that the effect was driven by increased efficiency within all task states compared with Pre-and Post-task resting states (Sensory: *z-range* = 2.4 – 4.69, *p <* 0.02, FPV: z-range = 3.78 – 5.22, p < 0.001).

**Figure 5.**
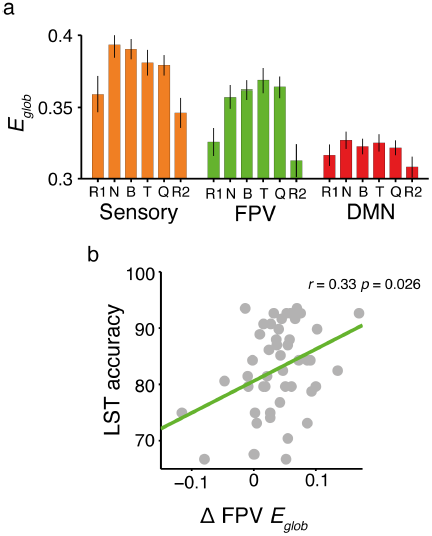
Changes in global network efficiency (E_glob_) across the identified reasoning task modules. **a.** Global network efficiency levels within each module across experiment conditions. Error bars represent 95% confidence intervals. R1= Pre-task rest, B = Binary, T = Ternary, Q = Quaternary, R2 = Post-task rest. **b.** Correlation between accuracy in the Latin Square Task (LST) and changes in fronto-parietal-visual (FPV) module efficiency during the task. Changes in network efficiency were correlated with overall reasoning performance, such that increased efficiency correlated with better task performance (*r* = 0.33, *p* < 0.01). Results are visualized at 15% network density.

We also investigated the relationship between individual differences in module efficiency and behavioral performance. To do so, we correlated reasoning accuracy scores with changes in module efficiency between the Pre-task resting state and the most complex reasoning condition (Quaternary). Only module efficiency within the FPV module was significantly correlated with behavior (see **Figure 5b**, mean statistics across thresholds: r = 0.33, p = 0.026; Spearman’s rs = 0.27, p = 0.084), such that larger increases in efficiency within the FPV module were associated with better reasoning performance. Neither the Default-mode or Sensory modules demonstrated such a relationship (p = 0.19, p = 0.29 respectively). Further, to probe the reliability of the above finding, we compared change in efficiency from Quaternary to the Null-task state, which yielded a similar result (r = 0.35, p = 0.021, Spearman’s rs = 0.30, p = 0.048). The correlation was also robust to partialing out fluid intelligence scores based on the Raven’s Matrices test (r = 0.35, p = 0.025). By contrast, there was no correlation between performance in the visual search task and module efficiency (N = 43, p = 0.65), suggesting the module efficiency-behavior relationships were specific to the LST. Finally, we also replicated these results, as well as the follow up Variation of Information results (**Figure 3d**), using the original visual, sensory, default-mode and fronto-parietal networks defined by Power and colleagues (2011), instead of our own data-driven modules.

## Discussion

Human reasoning has a quantifiable capacity limit (Halford et al., 1998). This limit is thought to arise from the brain’s ability to reconfigure interactions between spatially distributed networks (Cocchi et al., 2014; Parkin et al., 2015; Schultz and Cole, 2016), but recent work has highlighted the circumscribed nature of such interactions when compared with whole brain ‘resting’ architecture (Cole et al., 2014). In light of these recent findings, we examined how global and selective network properties change from resting to reasoning states, and how such changes relate to reasoning behavior. We found that complexity-based limits in reasoning ability rely on selective patterns of connectivity that emerge in addition to a more general task-induced functional architecture.

We used a non-verbal reasoning task, originally designed to test predictions from *relational complexity theory,* to systematically manipulate reasoning complexity (Halford et al., 1998; Birney et al., 2006; Birney and Bowman, 2009). In doing so we replicated previous behavioral results by demonstrating a reliable reduction in accuracy and an increase in reaction time as a function of increased complexity (Birney et al., 2006; Zhang et al., 2009; Zeuch et al., 2011). Importantly, an analysis of participants’ trial-by-trial ratings indicated that task errors were related to complexity demands and not factors such as transitory lapses in attention or disengagement from the task. The behavioral results also confirmed previous reports that individual reasoning capacity limits are correlated with scores on standard measures of fluid intelligence (Birney et al., 2006; Bhandari and Duncan, 2014) such as Raven’s Matrices.

Parametric increases in relational complexity have previously been tied to neural activity of segregated regions of the prefrontal cortices (Christoff et al., 2001; Kroger, 2002; Bunge et al., 2009; Golde et al., 2010) as well as to functional connectivity within fronto-parietal and cingulo-opercular “multiple-demand” networks (Cocchi et al., 2014; Parkin et al., 2015; Crittenden et al., 2016). Cingulo-opercular connectivity has been associated with initiating and maintaining task sets (Dosenbach et al., 2006) whereas the fronto-parietal network has been associated with moment-to-moment cognitive control (Cole and Schneider, 2007). Here we found that reasoning performance was best explained by a module composed of brain regions within the fronto-parietal, salience, subcortical and visual networks. First, functional connectivity and global efficiency of this subnetwork increased in line with increased reasoning demands (**Figures 4 & 5**). Second, larger increases in global efficiency within this module were associated with higher accuracy in the reasoning task. Finally, edge-wise connectivity of the fronto-parietal network was shown to increase in line with relational complexity, largely between default-mode and subcortical networks. These findings are broadly consistent with previous work showing that enhanced global network efficiency, and connectivity within the default-mode and fronto-parietal networks at rest, can predict intelligence and reasoning performance (Song et al., 2008, 2009; van den Heuvel et al., 2009; Finn et al., 2015; Hearne et al., 2016). Taken together the results confirm the central role of flexible fronto-parietal connectivity in implementing external goal-directed cognitive control (Cole et al., 2013, Cocchi et al., 2014).

It has been proposed that the cingulo-opercular network can be further divided to include a separate ‘salience’ system associated with bottom-up attention (Seeley et al., 2007; Power et al., 2011). Here we found that the salience network was implicated in the fronto-parietal-visual module but did not show complexity-induced edge-wise connectivity changes. On the other hand, the cingulo-opercular network did show edge-wise connectivity changes in line with reasoning complexity, but was not implicated in the fronto-parietal-visual module. This set of results is consistent with the notion that the cingulo-opercular network is a control-related counterpart of the fronto-parietal network, and suggests that the cingulo-opercular network might have a distinct role from that of the salience aspect of the system (Power et al., 2011).

It remains unclear precisely how ‘resting state’ networks coordinate flexible patterns of integration and segregation as a function of task complexity. Nevertheless, our findings support a key role for subcortical structures such as the thalamus in mediating such relationships (Bell et al., 2016; Sherman, 2016). Specifically, bilateral putamen and thalamus were implicated in both subnetworks that increased and decreased functional connectivity as task complexity increased (general trend shown in **Figure 4b**). This finding is in line with recent descriptions of the thalamus as a ‘global kinless’ hub, with evidence of activation in multiple cognitive contexts and strong connectivity across multiple large-scale functional networks (Guimera et al., 2007, Hwang et al., 2017, van den Heuval & Sporns, 2011). Such subcortical regions might be interpreted as managing relationships between task-related networks to form a coherent modular structure. Further work will be needed to elucidate the particular role of subcortical regions in this relatively unexplored area (Bell & Shine, 2016).

Functional brain module reconfigurations have previously been related to performance on a range of higher cognitive tasks, including learning (Bassett et al., 2011), working memory (Braun et al., 2015; Vatansever et al., 2015, 2017) and cognitive control (Dwyer et al., 2014). Here we found that the community architecture of the brain is flexible, but only in response to large cognitive shifts. For example, resting state visual and fronto-parietal modules, each of which is composed of several known sub-networks (Power et al., 2011), merged together during the reasoning task (**Figure 3**). Importantly, this reorganization was relatively isolated; follow up analyses indicated that the rest of the brain remained stable across changes in task complexity. In line with this observation, recent network-based re-conceptualizations of global workspace theory (Dehaene et al., 1998; Kitzbichler et al., 2011) have suggested that large, task-based module reconfigurations arise to better serve network communication underpinning behavior. Our work refines this idea by showing that once resting state modules reconfigure in response to external task demands, the majority of connectivity changes occur without interrupting the newly established modular architecture, as illustrated schematically in **Figure 6**. Moreover, it is these connectivity changes *within* the newly reconfigured modules that seem to be most related to behavior.

**Figure 6.**
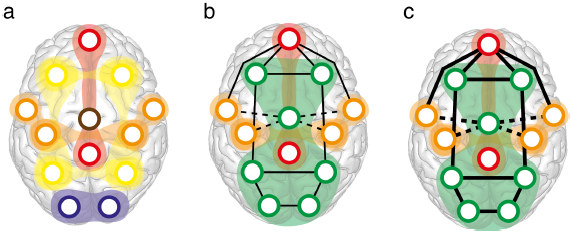
Conceptual model of functional networks supporting reasoning and rest states. **a.** At rest, functional modules are relatively independent. **b.** External, goal-directed task states are accompanied by broad module-level changes; a fronto-parietal-visual module forms (green), amongst stable default-mode (red) and sensory-motor modules (orange). **c**. Increased task demands are accompanied by specific increases (solid lines) and decreases (dashed-lines) in functional connectivity, rather than further modular reconfiguration. Ultimately, in the most complex conditions, the entire network reaches a similar level of correlation through both integrated and segregated dynamics (see **Figure 4b**).

Our finding that increased demands on cognitive reasoning are paralleled by a reduction in network modularity and increased efficiency has now been reported in several different task contexts (Kitzbichler et al., 2011; Bola and Sabel, 2015; Godwin et al., 2015; Vatansever et al., 2015; Cohen and D’Esposito, 2016; Shine et al., 2016; Westphal et al., 2017; Yue et al., 2017). Using the network-based statistic we found that these global changes were supported by increases and decreases in functional connectivity across multiple large-scale networks. Interestingly, the default-mode network has been suggested to act as a ’global integrator’, facilitating fronto-parietal cognitive control networks during conscious processing of information (Dehaene et al., 1998; Guldenmund et al., 2012; Leech et al., 2012; Vatansever et al., 2015). In line with this notion, we found that default-mode regions such as medial frontal cortex, angular gyri and posterior cingulate cortex demonstrated increased functional connectivity with fronto-parietal, cingulo-opercular and visual networks as task demands increased.

Modulations of visual network connectivity might be related to the visual nature of the LST, and specifically the requirement that participants search the 4 × 4 matrix to identify a shape at the probed location. A previous behavioural study found that participants’ eye fixation patterns differed for one-object and two-object relational problems (Gordon & Moser, 2007), raising the possibility that changes in relational complexity might be associated with changes in search patterns (and by extension, associated network connectivity). We cannot unequivocally rule out potentially small differences in eye movement patterns between complexity conditions in our study, but there are at least two reasons why such findings are unlikely to be directly relevant here. First, Gordon and Moser (2007) actively encouraged visual search by having their participants compare two different picture stimuli arranged one above the other on a page. By contrast, our LST paradigm involved the relatively brief presentation of a single 4 x 4 matrix at fixation. Second, the stimuli used by Gordon and Moser (2007) were visually complex line drawings that included several different object types that varied in size, shape and semantic content across trials. By contrast, our LST stimuli involved a single matrix containing identical shapes across complexity conditions, and did not explicitly require active search to solve for the target. Finally, we found no correlation between performance on a standard visual search task and reasoning performance or brain-based network efficiency metrics (subset of participants, N=43). If increasing visual search demands across task conditions was responsible for the observed network differences, performance on this visual search task should have correlated with the brain-derived metrics.

In conclusion, our findings suggest that reasoning demands rely on selective patterns of connectivity within fronto-parietal, salience, cingulo-opercular, subcortical and default-mode networks, which emerge in addition to a more general, task-induced modular architecture. Further work will be needed to elucidate the network processes that bring about the intricate and coordinated changes in connectivity patterns at the level of edges, modules and the whole brain, in the service higher cognition. Meanwhile, the current results provide novel insights into the roles of both specific and global network changes in reasoning.

## Acknowledgments

This research was supported by the Australian Research Council (ARC) ARC-SRI Science of Learning Research Centre (SR120300015), and the ARC Centre of Excellence for Integrative Brain Function (ARC Centre Grant CE140100007). J.B.M was supported by an ARC Australian Laureate Fellowship (FL110100103). L.C. was supported by an NHMRC grant (APP1099082). A.Z. was supported by a NHMRC Career Development Fellowship (GNT1047648). L.J.H was supported by an Australian Postgraduate Award. The authors declare no competing financial interests. We thank Oscar Jacoby and Zoie Nott for data collection assistance, Tong Wu for assistance with imaging analysis and Dr Kieran O’Brien, A/Prof Markus Barth and Dr Steffen Bollmann for performing the sequence optimizations.

